# Replacement of Arabidopsis H2A.Z with human H2A.Z orthologs reveals extensive functional conservation and limited importance of the N-terminal tail sequence for Arabidopsis development

**DOI:** 10.1101/2023.11.03.565555

**Authors:** Paja Sijacic, Dylan H. Holder, Ellen G. Krall, Courtney G. Willett, Maryam Foroozani, Roger B. Deal

**Author notes:** Correspondence: Roger B. Deal.

## Abstract

The incorporation of histone variants, distinct paralogs of core histones, into chromatin affects all DNA-templated processes in the cell, including the regulation of transcription. In recent years, much research has been focused on H2A.Z, an evolutionarily conserved H2A variant found in all eukaryotes. In order to investigate the functional conservation of H2A.Z histones during eukaryotic evolution we transformed *h2a.z* deficient plants with each of the three human H2A.Z variants to assess their ability to rescue the mutant defects. We discovered that human H2A.Z.1 and H2A.Z.2.1 fully complement the phenotypic abnormalities of *h2a.z* plants despite significant divergence in the N-terminal tail sequences of Arabidopsis and human H2A.Zs. In contrast, the brain-specific splice variant H2A.Z.2.2 has a dominant-negative effect in wild-type plants, mimicking an H2A.Z deficiency phenotype. Furthermore, H2A.Z.1 almost completely re-establishes normal H2A.Z chromatin occupancy in *h2a.z* plants and restores the expression of more than 84% of misexpressed genes. Finally, we used a series of N-terminal tail truncations of Arabidopsis HTA11 to reveal that the N-terminal tail of Arabidopsis H2A.Z is not necessary for normal plant development but does play an important role in mounting proper environmental stress responses.

**Article Summary:** H2A.Z is a histone variant conserved across the eukaryotes that plays important roles in transcription. Despite high overall conservation of H2A.Z protein sequence between organisms, sequence variation is seen in the N-terminal region, a site for many posttranslational modifications implicated in H2A.Z function. We replaced *Arabidopsis thaliana* H2A.Z with each of the human H2A.Z orthologs and found that both human H2A.Z isoforms rescue the severe phenotypes of plant H2A.Z null mutants, despite divergent N-terminal sequences. We further analyzed N-terminal sequence requirements by progressively deleting the endogenous plant H2A.Z N-terminus. Plants with a tailless version of H2A.Z developed normally but were impaired in stress responsiveness. Our results indicate that human H2A.Zs are broadly functional in plants and that the N-terminus of H2A.Z is not essential for normal development but is implicated in stress responsiveness.

## INTRODUCTION

Nucleosomes, the basic units of chromatin organization, are comprised of approximately 147 bp of DNA wrapped around a histone octamer, which contains the two copies of each of the four canonical histones H2A, H2B, H3, and H4. While the core histones are synthesized during the S-phase of the cell cycle for deposition onto a newly replicated DNA molecule, their counterparts, histone variants, are expressed throughout the cell cycle and can replace evicted canonical histones in a replication-independent manner [1–3]. The incorporation of histone variants has a profound effect on nucleosome function and affects virtually all DNA-templated cellular processes, including transcription [2–4].

H2A variants represent the largest and most diverse family of histones [2, 4, 5]. One member of this family is H2A.Z, a highly conserved variant of canonical H2A histone found in all eukaryotic organisms. Even though H2A.Z shares about 60% amino acid identity with canonical H2A [6, 7], the primary sequence differences between these two histones, particularly in the C-terminus of the proteins involved in intra- and internucleosomal interactions, led researchers to hypothesize that incorporation of H2A.Z into chromatin could alter the stability of nucleosomes and affect DNA folding [7, 8]. Studies have revealed H2A.Z to be involved in many processes including DNA repair and maintenance of genome stability, heterochromatin formation, telomere silencing, and both positive and negative regulation of transcription in animals and plants [9–16].

In the human genome, two genes encode H2A.Z proteins: *Homo Sapiens H2A.Z.1* and *H2A.Z.2.1* (*HsH2A.Z.1* and *HsH2A.Z.2.1*). Mammalian H2A.Zs are necessary for proper development as null mutations are embryonic-lethal [17]. HsH2A.Z.1 and HsH2A.Z.2.1 differ in only three amino acids, indicating a high level of functional redundancy. Interestingly, experimental evidence also suggests distinct functional roles for each HsH2A.Z protein [4, 16, 18, 19]. Recently, an alternatively spliced form of HsH2A.Z.2.1, named HsH2A.Z.2.2, was discovered and found to be predominantly expressed in brain tissues [20, 21]. Thus, humans possess three functional H2A.Z proteins with both redundant and specific roles during development.

In *Arabidopsis thaliana*, three genes encode H2A.Z proteins: *AtHTA8*, *AtHTA9*, and *AtHTA11*. Arabidopsis H2A.Z proteins act redundantly since mutations in the two most highly expressed H2A.Z genes (*AtHTA9* and *AtHTA11*) are necessary to detect pleiotropic morphological abnormalities. *H2A.Z* mutant plants experience a variety of phenotypic defects, including early flowering, lack of shoot apical dominance, altered flower development and reduced fertility, serrated leaves, and inability to respond to various biotic and abiotic stresses [22–25]. However, unlike animals, Arabidopsis *h2a.z* mutants are viable and fertile.

Interestingly, Arabidopsis H2A.Z proteins share more amino acid identity with human H2A.Zs than with other Arabidopsis H2A histones, indicating a high degree of H2A.Z evolutionary conservation [14]. The majority of sequence differences between Arabidopsis and human H2A.Zs are found at the N-terminal end, and, to a lesser extent, at the very C-terminal end of the proteins. In humans, many amino acids at the N-terminus, including multiple lysine residues, are known substrates for post-translational modifications (PTMs), which are shown to play an important role in H2A.Z-mediated regulation of transcription [15, 26, 27]. For instance, H2A.Z acetylation of lysine residues has been positively correlated with gene activation in many studies [26, 28–30], while H2A.Z lysine methylation has been associated with both gene repression and gene induction [31, 32]. Considering that the highest sequence divergence between the human and Arabidopsis H2A.Z proteins is in the N-terminal tail, we wondered how functionally conserved human and plant H2A.Zs are.

Here, we address this question by first generating a complete *h2a.z* knockout in Arabidopsis using CRISPR methodology, followed by the transformation of *h2a.z* plants with each of the three human *H2A.Z* genes. We show that human H2A.Z.1 and H2A.Z.2.1 phenotypically fully rescue the severe pleiotropic defects of Arabidopsis *h2a.z* plants, while the brain-specific splice variant H2A.Z.2.2 fails to complement these defects and in fact has a dominant negative effect in wild-type (WT) plants. At the molecular level, human H2A.Z.1 occupied nearly 100% of normal Arabidopsis H2A.Z deposition sites. Out of 8,399 mis-expressed genes in *h2a.z* plants, human H2A.Z.1 restored the expression of more than 84% of those genes back to their WT levels. Taken together, these results indicated a high degree of functional conservation between Arabidopsis and human H2A.Zs. Furthermore, this finding also led us to hypothesize that the N-terminal tails of Arabidopsis H2A.Zs are not necessarily essential for normal Arabidopsis development. Surprisingly, this hypothesis was supported by the ability of various N-terminal end truncations of Arabidopsis HTA11 to rescue the *h2a.z* phenotypic defects under normal growth conditions. However, when the N-terminal tailless HTA11 transgenic plants were exposed to abiotic stresses they showed significant growth defects, suggesting that the N-terminal tail of the H2A.Z protein may play an important role in the rapid gene induction and repression events required during stress responses. Future experimental evaluation, including the full identification of post-translational modifications to Arabidopsis H2A.Zs, is necessary to further dissect the role of the N-terminal tail in H2A.Z-mediated control of gene expression.

## RESULTS

### Human H2A.Z.1 and H2A.Z.2.1, but not H2A.Z.2.2, completely rescue the severe phenotypic defects of *h2a.z* plants

Our phylogenetic analysis of the H2A family of proteins from Arabidopsis and humans, together with other studies, revealed an interesting phenomenon: amino acid sequences of Arabidopsis H2A.Z proteins, AtHTA8, AtHTA9, and AtHTA11, are more similar to human H2A.Zs than to other Arabidopsis HTA (core H2A) histones (Fig 1A, [7, 14]). In fact, Arabidopsis and human H2A.Zs share about 80% amino acid identity, with the most diverse amino acid sequences between the proteins residing in their N-terminal ends (Fig 1B). This observation suggested a high degree of evolutionary conservation between human and Arabidopsis H2A.Zs and raised an intriguing question whether human H2A.Zs can function in plants and rescue the Arabidopsis *h2a.z* mutant defects.

**Figure 1.**
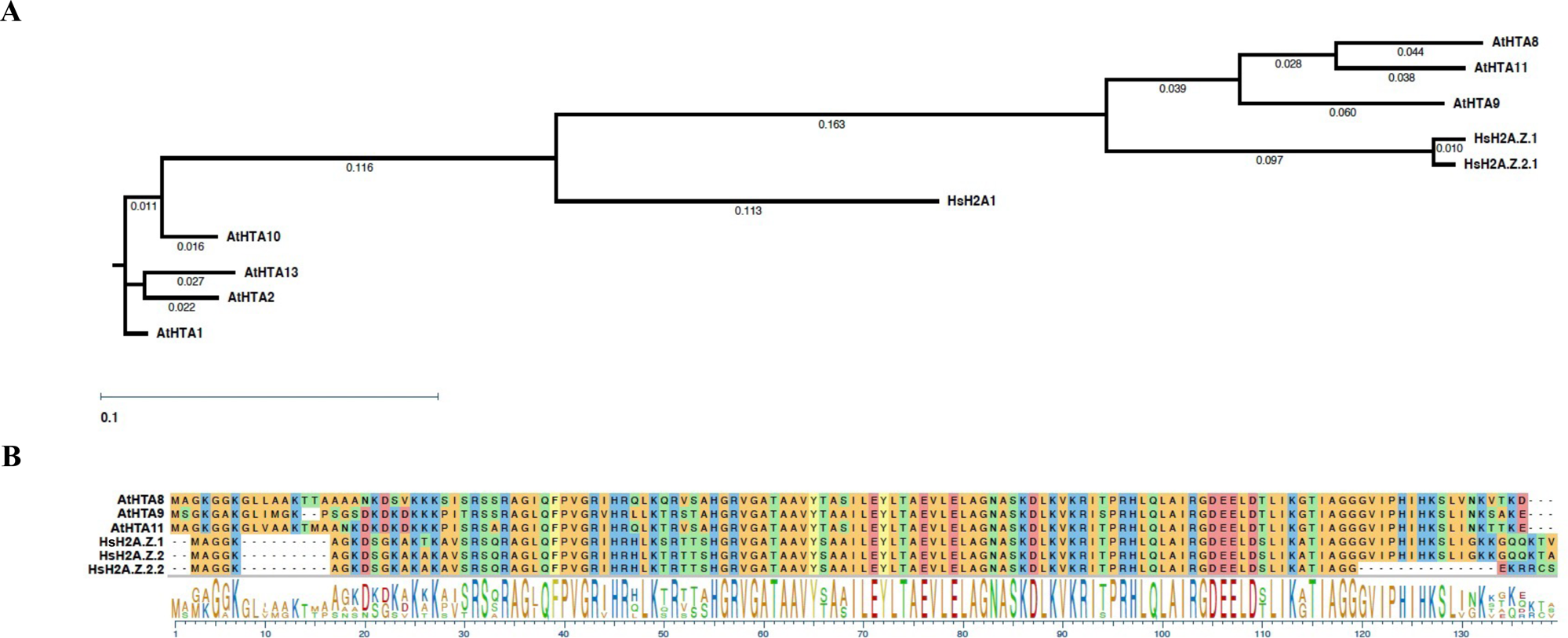
Arabidopsis H2A.Zs share about 80% amino acid identity with human H2A.Zs and are more similar to human H2A.Zs than to the other Arabidopsis H2A histones. **(A)** Phylogenetic tree of human and Arabidopsis H2A and H2A.Z proteins. **(B)** ClustalW alignment of three human H2A.Zs and three Arabidopsis H2A.Zs. Phylogenetic tree and sequence alignments were generated using DNASTAR.

To objectively assess the ability of human H2A.Z proteins to complement the phenotypic and molecular defects caused by the lack of endogenous H2A.Z in Arabidopsis we needed to use complete loss-of-function *h2a.z* plants. However, except for one example [33], all *h2a.z* mutant plants described in the literature are T-DNA-based double and triple insertion mutants, which are not complete *h2a.z* knockouts. Therefore, we first produced *h2a.z* null plants by employing the <u>C</u>lustered <u>R</u>egularly <u>I</u>nterspaced <u>S</u>hort <u>P</u>alindromic <u>R</u>epeats (CRISPR) methodology [34]. Arabidopsis plants identified as CRISPR-generated *h2a.z* homozygous triple knockouts (or simply *h2a.z* plants) have much more severe phenotypic defects than *h2a.z* T-DNA mutants, including drastically delayed germination, extremely stunted growth, pointy and twisted leaves, and complete sterility (Fig 2A, S1 Fig). In fact, very few of the *h2a.z* plants transitioned to flowering while the rest died prematurely. The *h2a.z* plants that do successfully transition to the reproductive stage of development have severely impaired flower development and are sterile (S1 Fig). We then identified the plants that are homozygous for CRISPR-generated mutant alleles of *hta9* and *hta11* and heterozygous for *hta8* CRISPR allele (hereafter named *h2a.z +/−* plants). These plants display phenotypic defects typical of T-DNA *h2a.z* mutants: serrated leaves, loss of apical dominance, aberrant petal number, partial fertility, and early flowering (Fig 2A, S1 and S2 Figs). Since *h2a.z +/−* plants are fertile, they were used for transformation with three plant codon-optimized human *H2A.Z* transgenes (*HsH2A.Z.1*, *HsH2A.Z.2.1*, and the *HsH2A.Z.2.1* splice variant known as *H2A.Z.2.2*). Importantly, we also included the endogenous genomic *HTA11* construct (*AtHTA11*) as our positive control for complementation, and a canonical H2A-encoding gene *HTA2*, as our negative control in rescue experiments. All constructs were driven by the native *HTA11* promoter, except for *HTA2*, which was driven by the constitutively active *35S* promoter. After the transformation of *h2a.z +/−* plants with different constructs, *h2a.*z null plants were identified among the progeny via genotyping. All subsequent assays were therefore performed on complete *h2a.z* null plants rescued with various constructs.

**Figure 2.**
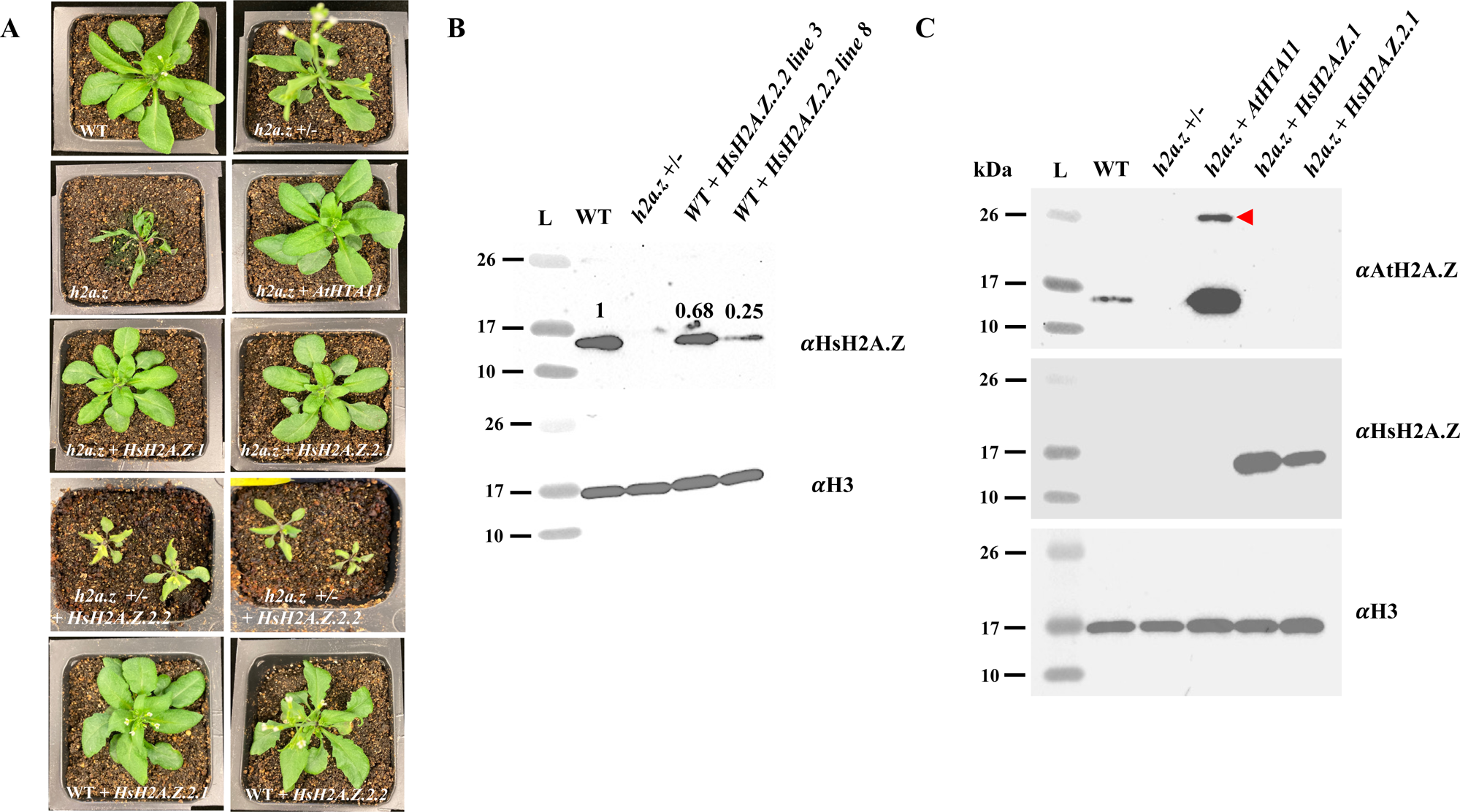
Human H2A.Z.1 and H2A.Z.2.1, but not H2A.Z.2.2, rescue phenotypic defects of crispr-generated *h2a.z* null plants. **(A)** 2.5 weeks old WT and mutant plants were grown under long-day conditions and individually photographed. **(B)** Western-blot analysis of leaf chromatin extracts from wild type (WT), *h2a.z +/−,* and two WT plants transformed with human HsH2A.Z.2.2, were probed with plant specific H2A.Z antibody (top panel) while H3 antibody was used as a loading control (bottom panel). The numbers above the H2A.Z western bands represent the ratio of signal intensities between the H2A.Z band and the corresponding H3 band, where the ratio for the WT plant is set to 1. The signal intensities were quantified using ImageJ. (**C**) Western-blot analysis of leaf chromatin extracts from wild type (WT), *h2a.z +/−,* and transgenic plants, were probed with either plant specific (top panel), or human-specific (middle panel) H2A.Z antibodies. H3 antibody was used as a loading control (bottom panel). The red arrowhead points to a 26 kDa monoubiquitinated form of Arabidopsis HTA11 detected in *h2a.z* + *AtHTA11* transgenic plants.

As expected, At*HTA11* was able to fully rescue all phenotypic defects of *h2a.z* plants (Fig 2A). Interestingly, *h2a.z* plants carrying either human *H2A.Z.1* or *H2A.Z.2.1* transgenes were also phenotypically rescued and indistinguishable from the WT and *h2a.*z + *AtHTA11* transgenic plants (Fig 2A, S2 Fig). On the other hand, T_1_ transgenic *h2a.z +/−* plants carrying a brain-specific *HsH2A.Z.2.2* variant had more severe defects than *h2a.z +/−* plants alone and more closely resembled *h2a.z* null phenotypes (Fig 2A). Moreover, 26 out of 29 T_1_ *HsH2A.Z.2.2* transgenic *h2a.z +/−* plants died before transitioning to the reproductive stage. Based on these results we hypothesized that the presence of human H2A.Z.2.2 in plants can somehow interfere with the normal function of endogenous Arabidopsis H2A.Zs. To test this possibility, we expressed HsH2A.Z.2.2 in WT plants and, as a control, we also expressed HsH2A.Z.2.1 in WT plants. We found that T_1_ plants carrying *HsH2A.Z.2.2* phenotypically resembled *h2a.z +/−* plants, including the early flowering phenotype, while the primary transgenic plants expressing *HsH2A.Z.2.1* were morphologically indistinguishable from WT plants (Fig 2A, S2 Fig). Furthermore, western analysis of two T_1_ WT plants carrying the *HsH2A.Z.2.2* transgene revealed that the incorporation of Arabidopsis H2A.Z proteins into chromatin was reduced to 68 and 25 percent, respectively, of the H2A.Z levels in untransformed WT plants (Fig 2B). These results support the notion that, when expressed in plants, human H2A.Z.2.2 can hinder the incorporation of endogenous Arabidopsis H2A.Zs.

On the other hand, T_1_ *h2a.z +/−* plants that expressed transgenic *HTA2* at high levels could not rescue the *h2a.z +/−* phenotypes (S3 Fig), suggesting that specifically only H2A.Z proteins, and not canonical H2A proteins, can complement the *h2a.z +/−* and *h2a.z* plant morphological defects. Overall, the results from our rescue experiments with human H2A.Zs suggest a high degree of functional conservation between Arabidopsis and human H2A.Z histones.

### Human H2A.Z.1 restores WT H2A.Z occupancy in *h2a.z* plants

Since both HsH2A.Z.1 and HsH2A.Z.2.1 equally complemented the phenotypic defects of *h2a.z* plants, we decided to use *h2a.*z + *HsH2A.Z.1* transgenic plants to examine the degree of molecular rescue in *h2a.*z plants mediated by human H2A.Z.1. To measure the ability of HsH2A.Z.1 to be deposited into chromatin, we performed two biological replicates of chromatin immunoprecipitation coupled with high-throughput sequencing (ChIP-seq) using a human-specific H2A.Z antibody on *h2a.*z + *HsH2A.Z.1* transgenic plants, and on WT and *h2a.*z + *AtHTA11* transgenic plants using an Arabidopsis-specific H2A.Z antibody (Fig 2C). When we compared the average patterns and heat maps of H2A.Z enrichment across genes in WT plants, *h2a.*z + *AtHTA11*, and in *h2a.*z + *HsH2A.Z.1* transgenic plants we found highly similar profiles, indicating that both *AtHTA11* and human *H2A.Z.1* are able to properly re-establish H2A.Z deposition sites in *h2a.z* plants (Fig 3A). We then compared H2A.Z ChIP-seq read counts on a per gene basis between WT and *h2a.*z + *AtHTA11* plants, and between WT and *h2a.*z + *HsH2A.Z.1* plants, and discovered that only four genes were differentially enriched for H2A.Z between WT and *h2a.*z + *AtHTA11*, and only 266 genes between WT and *h2a.*z + *HsH2A.Z.1* transgenic plants (Figs 3B-C). Overall, these results suggest that human H2A.Z.1 restores WT H2A.Z deposition sites in *h2a.z* plants almost completely.

**Figure 3.**
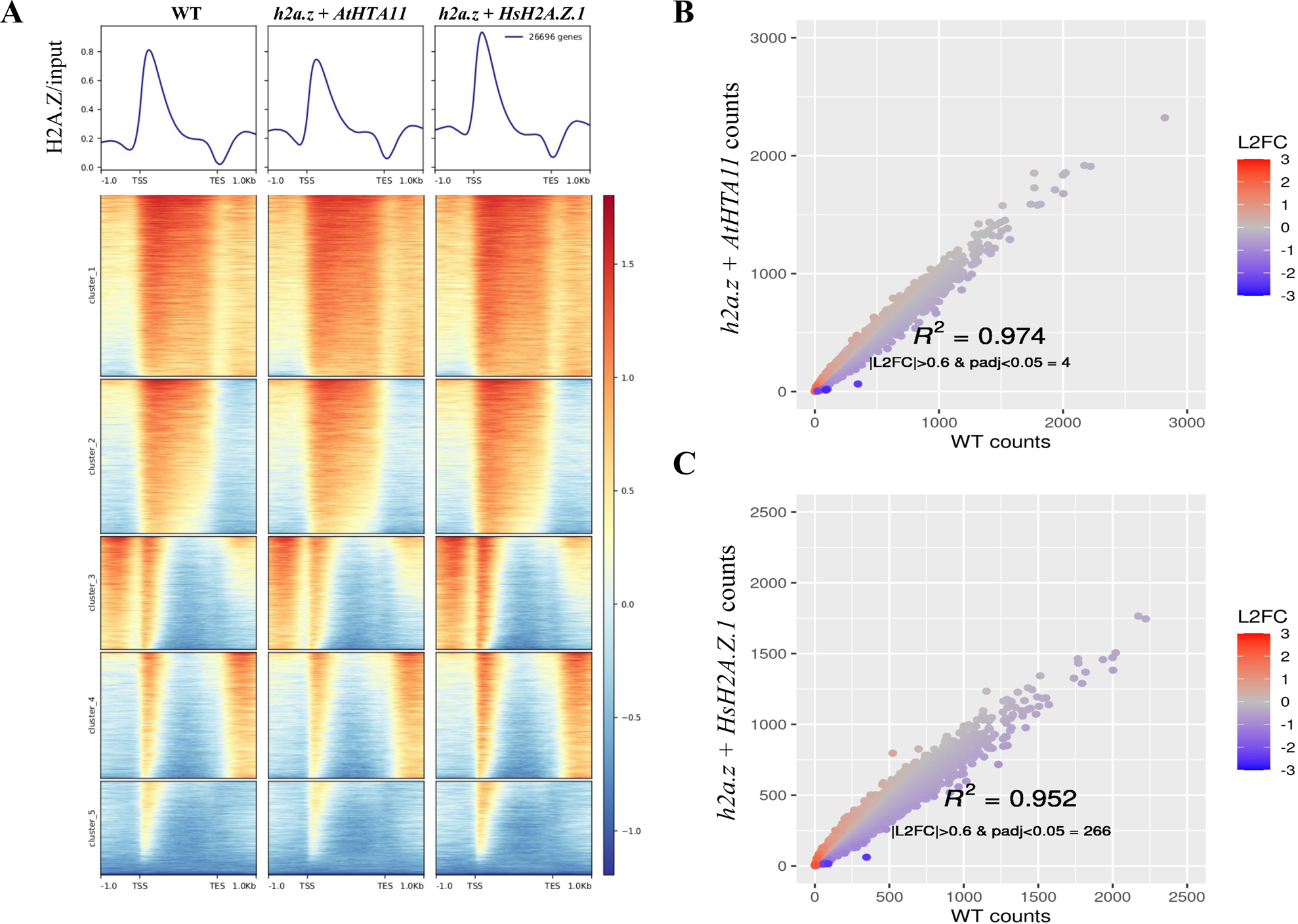
Chromatin immunoprecipitation with sequencing (ChIP-seq) analysis reveals that human H2A.Z.1 restores H2A.Z deposition sites in *h2a.z* plants. **(A)** Heatmap of normalized ChIP-seq signal (H2A.Z/input) for 26,669 genes. Average ChIP-seq H2A.Z profiles of WT, *h2a.z* + *AtHTA11*, and *h2a.z* + *HsH2A.Z.1* plants were plotted over gene body coordinates of 26,669 genes, from the transcript start site (TSS) to the transcript end site (TES) and +/− 1 kb from each end. Five k-means clusters were sorted by mean signal value. One replicate is used as a representative of each genotype. **(B and C)** Correlation of H2A.Z ChIP-seq read counts at genes between WT and *h2a.z* + *AtHTA11* plants (**B**), and between WT and *h2a.z* + *HsH2A.Z.1* plants (**C**). Read counts at genes were defined by mapped reads from the TSS to the TES. Counts were normalized via DESeq2. Color represents log2 fold change of normalized counts in sample vs WT.

### The expression of more than 84% of mis-expressed genes in *h2a.z* plants is restored to WT levels by human H2A.Z.1

After discovering near total restoration of H2A.Z deposition in *h2a.*z + *HsH2A.Z.1* plants, we performed RNA-seq experiments to investigate the ability of human H2A.Z.1 to restore the expression of misregulated genes in *h2a.z* plants. After isolating total RNA from three biological replicates of WT, *h2a.z* plants, *h2a.*z + *AtHTA11*, and *h2a.*z + *HsH2A.Z.1* transgenic plants we identified 8,399 differentially expressed genes (DEGs) in *h2a.z* plants when compared to the WT, with 3714 downregulated genes and 4685 upregulated genes (with p adj ≤ 0.05, and absolute log2 fold change ≥ 0.6, Figs 4A-B). A markedly lower number of genes were misexpressed in *h2a.*z + *AtHTA11* plants (809 downregulated genes and 137 upregulated genes, total of 946 DEGs), and in *h2a.*z + *HsH2A.Z.1* plants (1106 downregulated genes and 1332 upregulated genes, total of 2438 DEGs) when compared to the WT (Figs 4A-B). We then examined the overlap of all misexpressed genes in *h2a.*z plants with all misexpressed genes in *h2a.*z + *AtHTA11* plants and discovered that 593 genes, out of 8399, were still misregulated in *h2a.*z + *AtHTA11* plants (Fig 4C) while 353 genes were uniquely misexpressed only in *h2a.*z + *AtHTA11* plants. Overall, these results demonstrated that the *AtHTA11* transgene restores the expression of 93% of misexpressed genes in *h2az* plants.

**Figure 4.**
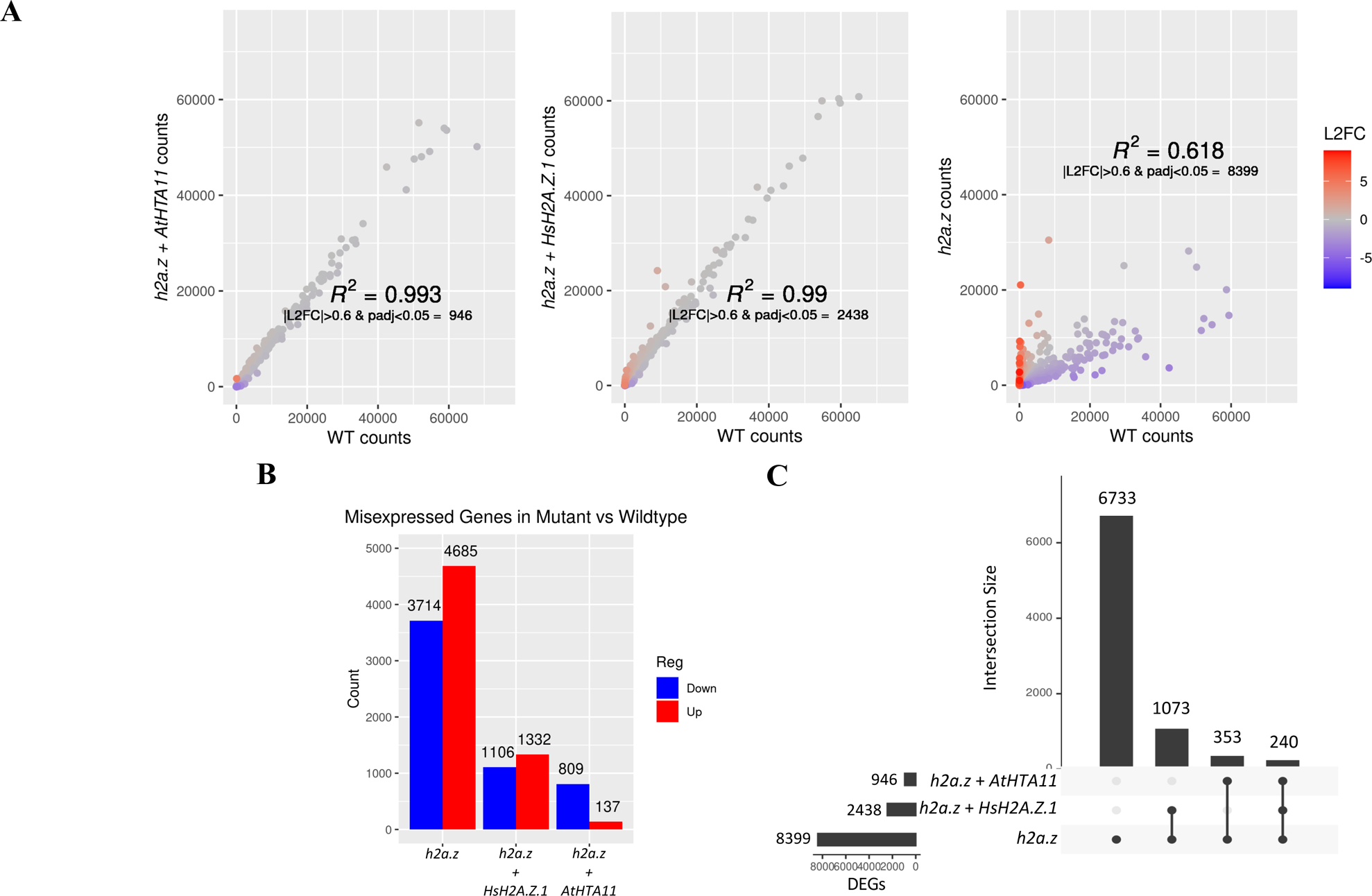
RNA-seq analysis reveals that human H2A.Z.1 restores the expression of more than 84% of misregulated genes in *h2a.z* plants back to WT levels. **(A)** Correlation of DESeq2 normalized transcript counts per gene between WT and *h2a.z* + *AtHTA11* plants (left), WT and *h2a.z* + *HsH2A.Z.1* plants (middle), and between WT and *h2a.z* plants (right). Color represents log2 fold change of normalized counts per gene in sample vs WT. **(B)** Histogram representing total number of misexpressed genes in mutant and transgenic plants versus WT. **(C)** Upset plot of shared differentially expressed genes across *h2a.z* + *AtHTA11*, *h2a.z* + *HsH2A.Z.1*, and *h2a.z* plants compared to WT. Differential expression is defined by |L2FC| > 0.6 and padj < 0.05.

Similarly, we analyzed the overlap of all misexpressed genes in *h2a.z* plants with all misexpressed genes in *h2a.*z + *HsH2A.Z.1* plants and found that 1,313 genes were still misregulated in *h2a.*z + *HsH2A.Z.1* plants (Fig 4C) while 1,125 genes were newly misexpressed only in *h2a.*z + *HsH2A.Z.1* plants. Based on these results we calculated that human H2A.Z.1 is able to restore the expression of more than 84% of *h2a.z* mis-expressed genes back to the WT levels.

Finally, we also examined the average H2A.Z enrichment in WT across the bodies of genes either downregulated or upregulated in *h2a.*z plants compared to WT and discovered that the genes upregulated in *h2a.z* have higher H2A.Z levels compared to downregulated genes (S4 Fig). These results are consistent with H2A.Z enrichment at gene bodies being negatively correlated with gene expression, as observed in previous studies [13]. Taken together, our results suggest that human H2A.Z.1 rescues the molecular defects of *h2a.z* plants at remarkably high level even though the amino acid sequences between Arabidopsis and human N-terminal and C-terminal ends are significantly different (Fig 1B).

### N-terminal truncated transgenes of Arabidopsis *HTA11* rescue majority of severe phenotypic defects of *h2a.z* plants

Because human H2A.Z.1 largely recapitulates the normal function of Arabidopsis H2A.Zs in transgenic plants despite the divergent N-terminal sequences between these proteins (Fig 1B), we hypothesized that post-translational modifications of amino acid residues in the N-terminal tail of Arabidopsis H2A.Zs are not critical for the overall function of H2A.Z in development. Since these PTMs are not well studied [35], instead of replacing the individual key residues within the N-terminus, we decided to employ a more drastic approach of testing this hypothesis by truncating the N-terminal tail of Arabidopsis HTA11 to various lengths. We engineered four different *AtHTA11* constructs, truncating 7, 13, 21, or 28 amino acids at the N-terminus, and transformed them into *h2a.z +/−* plants to assess their ability to rescue the mutant defects.

Phenotypic analysis of *h2a.z* plants carrying either of the four truncated *AtHTA11* constructs revealed that these plants closely resembled WT plants morphologically (Fig 5A) under normal growing conditions. We also found that the truncated AtHTA11 proteins are successfully deposited into chromatin (Fig 5B). Since these results were somewhat unexpected, especially for *AtHTA111128*, which is practically an N-terminal tailless H2A.Z protein, we wanted to assess if these plants could properly respond to stress conditions. To this end, we subjected *AtHTA111128* plants to two abiotic stresses: high salt stress and exogenous application of the plant phytohormone abscisic acid (ABA). When grown on ½ MS and 100mM NaCl plates (medium salt stress condition), both WT and *AtHTA11Δ28* plants were phenotypically indistinguishable from each other and germinated at similar rates (Fig 6A and S5A Fig). However, when these plants were exposed to high salt stress (200mM NaCl) *AtHTA11Δ28* transgenics germinated significantly less than WT plants, indicating that *Δ28 AtHTA11* plants are hypersensitive to high salt stress (Fig 6A, S5A Fig). Similarly, *AtHTA11Δ28* germinated at significantly lower rate than WT plants on ½ MS plates supplemented with 0.3 μM ABA, suggesting that *AtHTA11Δ28* plants are also more sensitive to ABA compared to WT plants (Fig 6B, S5B Fig). Overall, these results indicate that the N-terminal tail of H2A.Z is required to successfully mediate plant responses under the stress conditions tested. Based on these results we concluded that the N-terminal tail of Arabidopsis H2A.Z appears to be dispensable for its major functions during normal plant development but plays an important role in eliciting proper responses when plants are exposed to environmental stressors.

**Figure 5.**
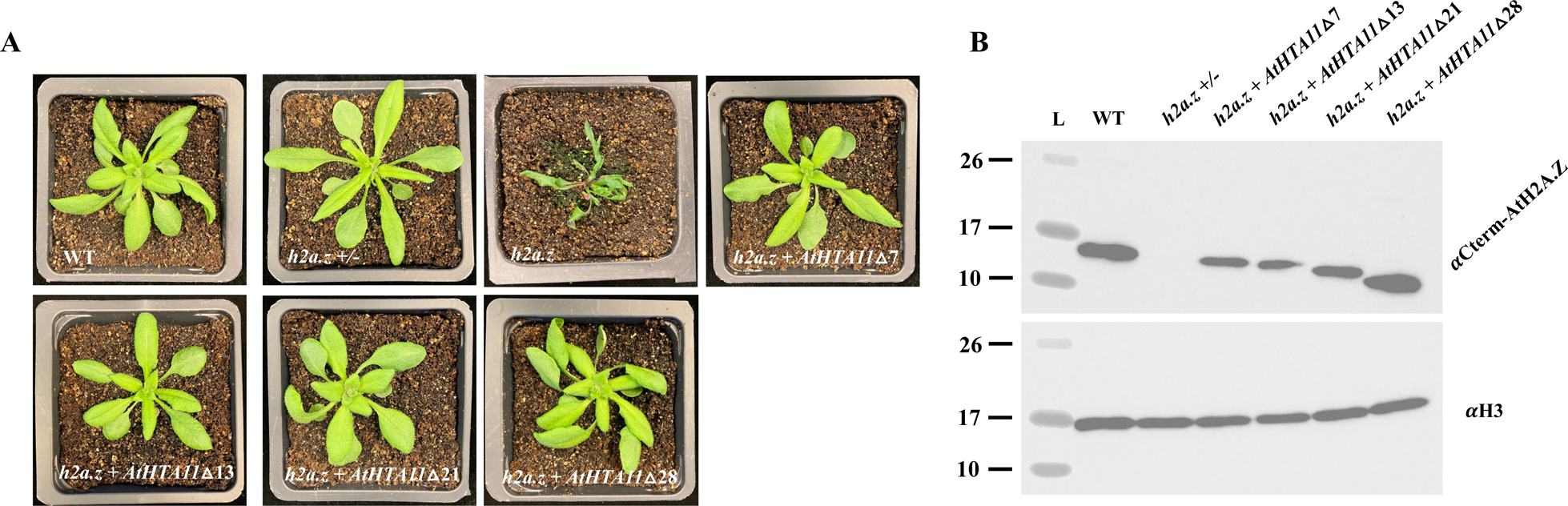
N-terminal truncations of AtHTA11 largely rescue *h2a.z* defects. **(A)** 2.5-week-old plants were grown under long-day conditions and individually photographed. **(B)** Western-blot analysis of leaf chromatin extracts from wild type (WT), *h2a.z* +/− plants, and T_2_ *h2a.z* plants with four different truncated Arabidopsis *HTA11* constructs probed with Arabidopsis H2A.Z specific antibody that recognizes the C-terminal end of the protein (top panel), and with an H3 antibody that was used as a loading control (bottom panel).

**Figure 6.**
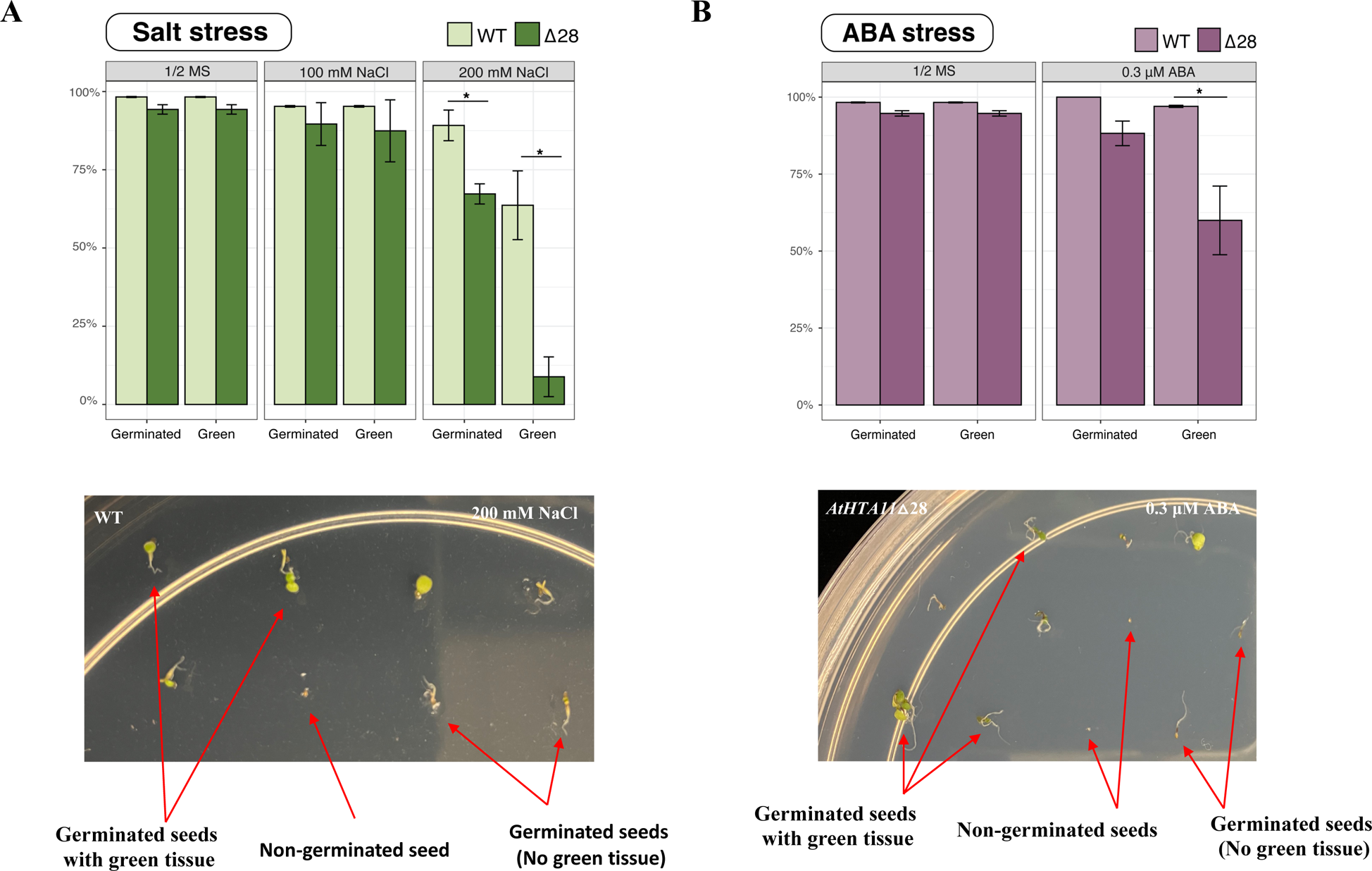
N-terminal truncated *AtHTA11*△28 plants are hypersensitive to high salt and ABA stress conditions when compared to WT plants. Eleven-day-old seedlings grown on ½ MS agar plates and plates supplemented with either 100 mM or 200 mM NaCl **(A)** or supplemented with 0.3 μM Abscisic acid (ABA) **(B)**, were characterized for their ability to germinate and to produce green tissue. **(A)** Germination rates for seeds undergoing medium salt stress (100 mM NaCl), and high salt stress (200mM NaCl) compared to normal growth conditions (½ MS). Germination rates from two independent germination experiments are stratified into two categories, “Germinated” which indicates any germinated seed with root or shoot tissue, and “Green” which indicates any germinated seed with green tissue (see bottom photo for phenotypic key). On ½ MS and medium salt stress, *AtHTA11*≥28 (≥28) and WT plants have equivalent germination percentages. At high salt stress, Δ28 plants have significantly less germination (p=0.034) and green tissue (p=0.026) when compared to WT plants (*student’s t-test, p<0.05). **(B)** Germination rates for plants undergoing Abscisic Acid (ABA) stress (0.3 μM) compared to normal growth conditions (½ MS). Germination rates from two independent germination experiments are stratified into two categories, “Germinated” which indicates any germinated seed with root or shoot tissue, and “Green” which indicates any germinated seed with green tissue (see bottom photo for phenotypic key). *AtHTA11*≥28 (≥28) and WT have equivalent germination rates on ½ MS media, and there is also no statistically significant difference in germination rate on ABA-containing media. However, Δ28 has significantly less germinated seeds with green tissue (p=0.042) on ABA-containing media compared to WT (*student’s t-test, p<0.05).

## DISCUSSION

### Canonical human H2A.Z proteins rescue morphological and molecular defects of *h2a.z* plants at a remarkably high level

We originally hypothesized that human H2A.Zs would not be able to rescue the major phenotypic defects of *h2a.z* plants since the functionally significant amino acid residues found at the N-terminal tails of these proteins are very different in sequence (Fig 1B). However, we found that human H2A.Z.1, when expressed in *h2a.z* plants, complemented not only the major morphological abnormalities of mutant plants but also rescued the molecular defects caused by the loss of endogenous H2A.Zs at a remarkably high level. On the other hand, human H2A.Z.2.2 had an opposite effect when expressed in plants and appears to interfere with normal H2A.Z deposition or retention in chromatin. Even though the mechanism of this obstruction is not known, human H2A.Z.2.2, in the form of an inducible construct, might be useful for temporally disrupting endogenous H2A.Z functions in plants.

Overall, these results clearly demonstrated that there is an incredibly high degree of functional conservation between Arabidopsis and human H2A.Z histones and raise an interesting question about the ability of Arabidopsis H2A.Zs to rescue any defects caused by the loss of H2A.Zs in human cells, or in other model organisms.

### The N-terminal tail of Arabidopsis H2A.Z is not required for normal development but plays an important role in proper environmental stress responses

To explain the finding that human H2A.Zs are highly functional in Arabidopsis, we contemplated two possible scenarios, which are not mutually exclusive: 1) Post-translational modifications (PTMs) of amino acids at the N-terminal end of Arabidopsis H2A.Zs are important for their overall function and plant enzyme machinery that deposits those modifications can correctly recognize key residues within the N-terminal tails of human H2A.Zs to properly modify them, 2) PTMs of the key amino acid residues at the N-terminal tail of Arabidopsis H2A.Zs are not important for their function and, therefore, human H2A.Zs may function in plants as efficiently as endogenous H2A.Zs regardless of their ability to be post-translationally modified by plant machinery.

Since PTMs of amino acids at the N-terminal tail of Arabidopsis H2A.Zs have not been well characterized [35], we utilized a more aggressive approach to test these hypotheses by transforming four different N-terminally truncated constructs of Arabidopsis *HTA11* into *h2a.z +/−* plants and assessing their ability to rescue the phenotypic defects of *h2az* null plants. Surprisingly, all four constructs successfully rescued *h2a.z* morphological defects, including *AtHTA111128*, indicating that the N-terminal tail of Arabidopsis H2A.Z is not essential for its function during normal development. At first, these results were quite unexpected. However, a recent study in budding yeast has demonstrated that it is not the N-terminal tail but rather the specific set of conserved amino acids at the C-terminus that contributes to the unique function of H2A.Z [36]. Indeed, all Arabidopsis and human H2A.Zs, except for a few residues that are missing in HsH2A.Z.2.2, possess these conserved amino acids, while they are absent from Arabidopsis HTA2 histone (S6 Fig).

It was difficult to imagine that the N-terminal tail, carrying many conserved amino acids known to be the substrates for PTMs, has no significance for H2A.Z biology in plants. For instance, Arabidopsis HTA11 contains 11 lysines at the N-terminus which are known targets of numerous modifications, and plants do possess enzymatic machinery capable of recognizing and modifying lysine residues [37–40]. Therefore, it is reasonable to assume that AtHTA11 lysine modifications would be required at some point during the plant life cycle. The question then arises, what role do modified amino acids at the N-terminus play in H2A.Z biology that we have not observed at the phenotypic level? One possibility is that the putatively modified residues may fine-tune H2A.Z function in response to stimuli, such as the exposure to abiotic stresses. We went on to test this hypothesis using *AtHTA11Δ28* plants and discovered that N-terminal tailless H2A.Z transgenic plants responded poorly to ABA and high salt stresses when compared to WT plants. These results suggest that the amino acids in the N-terminal tail of H2A.Z, likely with their corresponding post-translational modifications, may play an important role in properly mediating rapid transcriptional activation and repression events required during stress responses. Furthermore, our results also suggest that at least the C-terminal tails of human H2A.Z proteins are being properly post-translationally modified in Arabidopsis, as we have observed that HsH2A.Z.1 is apparently monoubiquitinated at the C-terminal end when expressed in plants (S7 Fig). However, to fully understand the potential roles of H2A.Z PTMs, we will need to comprehensively define them and to study how they change during transcriptional induction and repression events.

## MATERIALS and METHODS

### Plant material, growth conditions, and transformation

*Arabidopsis thaliana* of the Columbia (Col-0) ecotype was used as the wild-type reference. Seedlings were grown either in soil or on half-strength Murashige and Skoog (MS) media agar plates [41], in growth chambers at +20°C under a 16 hour-light/8-hour dark cycle with light intensity of 330 μmol/m^2^s. All plasmids used for plant transformation were introduced into *Agrobacterium tumefaciens* GV3101 strain by electroporation. T_0_ plants were transformed using the floral dip method [42]. Primary transgenic plants were selected on half-strength MS media agar plates containing 25 mg/L BASTA and 100 mg/L timentin, and then transferred to soil.

### Production of *h2a.z* plants using CRISPR

To generate CRISPR-induced *h2a.z* null mutant plants we utilized an egg cell-specific promoter-controlled Cas9 plasmid that contains two single guide RNAs, which can target two separate genes at the same time [43]. *HTA9* and *HTA11* were targeted first since *hta9/hta11* double mutants have easily observable phenotypes. We sequenced the regions around the target sites using DNA samples isolated from T_2_ transgenic plants with *h2a.z* phenotypes and confirmed that these plants were indeed double homozygous mutants for *hta9* and *hta11*. All mutations were simple base pair insertions causing a frame shift in the coding sequence leading to premature stop codons (S8 Fig). We then identified plants that were *hta9/hta11* null without the Cas9 transgene and transformed them with the same egg cell-specific promoter-controlled Cas9 plasmid, this time targeting the *HTA8* gene. After DNA regions around *HTA8* target sites were sequenced, we identified new primary transgenic plants that were homozygous for *hta9 and hta11* and heterozygous for *HTA8*, herein named *h2a.z +/−* plants. The *HTA8* CRISPR allele was a simple base pair insertion causing a frame shift in the coding sequence (S8 Fig). We then identified *h2a.z +/−* plants that segregated out the *Cas9* transgene and these were used for all transformation experiments. Eventually, we identified *h2a.z* null mutants among *h2a.z +/−* progeny plants and confirmed by Sanger sequencing that these plants indeed had mutations in all three *H2A.Z* genes.

### Plasmid DNA constructs

To produce all constructs that were driven by the native *HTA11* promoter we first cloned 1855 bp of the *AtHTA11* promoter sequence (1855 bp upstream from the *AtHTA11* ATG start codon), with added *NcoI* restriction site at the 3’ end, into gateway-compatible pENTR-D/TOPO plasmid (Invitrogen). We then cloned 482 bp of the *AtHTA11* terminator sequence (482 bp downstream from the *AtHTA11* stop codon) into the same pENTR plasmid that contained the *AtHTA11* promoter just of the *NcoI* restriction site. Various PCR-generated *AtHTA11* constructs and synthetic plant codon-optimized human *H2A.Zs* were then cloned into this plasmid using the *NcoI* restriction site. All constructs were verified by sequencing and eventually subcloned into pMDC123 gateway destination plasmid [44] using the LR clonase II enzyme in LR recombination reaction (Invitrogen). To produce *35S::HTA2* construct we first cloned *HTA2* gene sequence into pENTR-D/TOPO plasmid (Invitrogen). After the correct sequence of HTA2 was verified by Sanger sequencing, the construct was subcloned into pEG100 gateway destination plasmid [45] using the LR clonase II enzyme in LR recombination reaction (Invitrogen).

### cDNA production and real-time RT-PCR (qRT-PCR)

Total RNA was isolated from third and fourth pair of leaves from *35S::HTA2* transgenic *h2a.z +/−* plants using the RNeasy plant mini kit (Qiagen). 2 μg of total RNA was converted into cDNA with LunaScript RT SuperMix kit (New England Biolabs). The cDNAs were used as templates for real-time PCR and ran on StepOnePlus real-time PCR system (Applied Biosystems) using SYBR Green as a detection reagent. The *PP2A* gene (AT1G13320) was used as the endogenous gene expression control [46], while primers specific for the *35S::HTA2* transgene were used to detect its expression in three individual T_1_ *h2a.z +/−* plants.

### Chromatin extraction and protein gel blotting

To probe chromatin-associated proteins by western blotting we used the chromatin extraction protocol described by Luo and colleagues [47], with following modifications: 1) 0.5 grams of rosette leaves from each sample were ground in liquid nitrogen and homogenized in 5 ml of Honda buffer, 2) Homogenates were filtered through 70 μm cell strainer and centrifuged at 1,500g at + 4°C degree for 20 minutes. The chromatin fractions from each sample were resuspended in 80-100 μl of 1x Laemmli’s sample buffer. Western blotting was performed as previously described [48], using 1:1000 dilutions of following primary antibodies: N-terminal Arabidopsis H2A.Z antibody [22], affinity-purified rabbit polyclonal C-terminal Arabidopsis H2A.Z antibody (raised against the peptide sequence: DTLIKGTIAGGGVIPHIHKSLI), human H2A.Z antibody (Abcam, ab188314), and H3 antibody (Abcam, ab1791). All blots were incubated with ECL detection reagents for 2 minutes (Thermo Scientific) and scanned for chemiluminescence signal using ChemiDoc MP imaging system instrument (BioRad).

### Chromatin immunoprecipitation (ChIP) with Arabidopsis and human H2A.Z antibodies

ChIP experiments were performed in biological duplicates on WT, *h2a.*z + *AtHTA11*, and on *h2a.*z + *HsH2A.Z.1* leaf tissue (third and/or fourth pair of rosette leaves) as described previously [49], with following modifications: 1) For each sample, 0.5 grams of leaf tissue were used, 2) ground samples were lysed in 600 μl of buffer S, 3) around 400 μl of the slurry were transferred to 0.6 ml tubes and sonicated in the Bioruptor (Diagenode) at + 4°C for 1 hour on the “high” setting with sonication intervals set to 45 seconds on/15 seconds off, 4) The sonicated lysates were centrifuged at 20,000xg for 10 min at + 4°C and 300 μl of the supernatants were transferred to a fresh 5 ml tube where 2.7 ml of buffer F was added and mixed well, 5) 50 μl of this mixture was saved as the input sample while 1.5 ml was used for immunoprecipitation with specific antibodies, 6) Arabidopsis-specific H2A.Z antibody [22] and human-specific H2A.Z antibody (Abcam, ab188314) were added to the diluted lysates at a final concentration of ∼2 μg/ml and incubated overnight at + 4°C with rocking, 7) 25 μl of washed Protein A dynabeads were added to each sample and incubated at + 4°C for 2 hours with rocking, 8) immunoprecipitated DNA fragments were purified using Qiagen Minelute kit and eluted in 14 μl of elution buffer. Eluted DNA samples were quantified using Picogreen reagent (Thermo Fisher).

### ChIP-seq library preparation, sequencing, and data analysis

ChIP-seq libraries were prepared starting with ∼1000 picograms of ChIP or input DNA samples using the ThruPlex DNA-seq kit (Takara) according to the manufacturer’s instructions. Libraries were pooled together and sequenced using paired-end 150 nt reads on an Illumina NovaSeq 6000 instrument. Reads were trimmed of adapter content using Trimgalore [50] and mapped to the *Arabidopsis thaliana* Col-PEK genome assembly [51] using Bowtie2 [52] with the following parameters: --local --very-sensitive --no-mixed --no-discordant --phred33 -I 10 -X 700. Aligned reads were converted to the BAM file format and quality filtered using Samtools [53]. Duplicate reads were removed using Picard markDuplicates [54]. For visualization, deduplicated BAM files of each genotype were converted to bedgraph files using bedtools bamtobed followed by genomecov [55]. Samples were then normalized to the lowest read depth and converted to BigWig using bedGraphToBigWig [56]. Finally, the signal from each sample was expressed as the log2 ratio relative to the corresponding input using bigWigCompare with –psuedocount = 1 to avoid 0/x [57]. Profile plots and heatmaps of the resulting bigwig files were generated using Deeptools [57]. Mapped fragments overlapping with genes or peaks were counted using featureCounts [58]. Gene counts were then analyzed for differential enrichment using DESeq2 [59]. DESeq2 results were visualized using ggplot2 package in R [60].

### RNA extraction, RNA-seq library preparation, sequencing, and data analysis

Total RNA was isolated from the following plants: WT, *h2a.z*, *h2a.*z + *AtHTA11*, and *h2a.*z + *HsH2A.Z.1*. Five individual leaves (third or fourth pair of rosette leaves) were collected from 5 different plants of WT, *h2a.*z + *AtHTA11*, and *h2a.*z + *HsH2A.Z.1* genotypes. The leaves were ground in liquid nitrogen and powder was resuspended in 2 ml of RLT buffer from RNeasy Plant Mini kit (Qiagen). For each sample, three equal aliquots of 400 μl of this resuspension were further processed to extract total RNA, according to the manufacturer’s instructions. For *h2a.z* plants, aboveground tissue from three different plants was harvested and individually processed to extract total RNA following the manufacturer’s recommendations. Total RNA was DNase-treated (Ambion) and quantified using Nanodrop One (Thermo Scientific). 100 nanograms for each of the three replicates from every sample were then used as a starting material to generate RNA-seq libraries following the Universal RNA-seq with NuQuant protocol (Tecan). Libraries were pooled together and sequenced using paired-end 150 nt reads on an Illumina NovaSeq 6000 instrument. Reads were trimmed of adapter content using Trimgalore [50]. Trimmed reads were mapped to the *Arabidopsis thaliana* TAIR10 genome assembly, converted to the BAM file format, and sorted by coordinate using STAR [61]. Aligned reads were indexed and quality filtered using Samtools [53]. Mapped fragments overlapping with exons were counted using featureCounts [58]. Gene counts were then analyzed for differential enrichment using DESeq2 [59]. Significantly differentially expressed genes were defined as those having an adjusted p-value less than 0.05 and an absolute fold change greater than 1.5. DESeq2 results were visualized using ggplot2 and upsetR in R [60].

### Abiotic stress experiments

WT and *AtHTA111128* seedlings were grown on half-strength Murashige and Skoog (MS) media agar plates, or plates supplemented with either NaCl (100 mM or 200 mM) or with 0.3 μM ABA, in growth chambers at +20°C under a 16-hour light/8-hour dark cycle. Eleven-day-old seedlings were then photographed and characterized for their ability to germinate and to produce green tissue.

### Data availability

All sequencing data are available in the NCBI GEO database under accession number GSE263313.

## Acknowledgements

We thank members of the Deal lab for critical comments and suggestions on the manuscript.

---

**S1 Figure.**
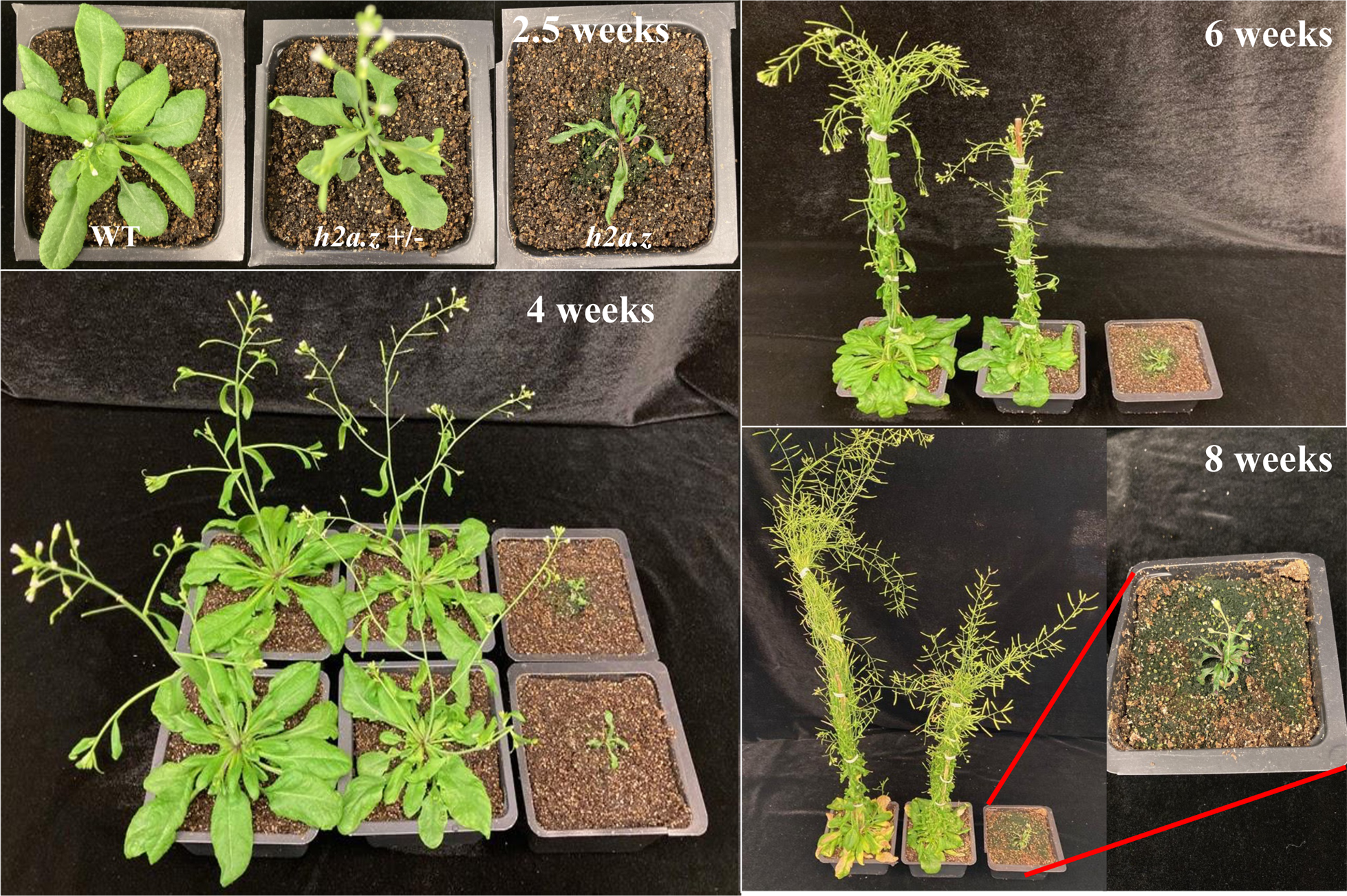
Complete loss-of-function *h2a.z* mutant plants have severe developmental defects. WT (left), *h2a.z* +/− (middle), and *h2a.z* (right) plants, grown under long-day conditions, were individually photographed at four time points over eight weeks of growth. *h2a.z* mutant plants are dwarfed and have severely delayed development compared to *h2a.z* +/− and WT plants.

**S2 Figure.**
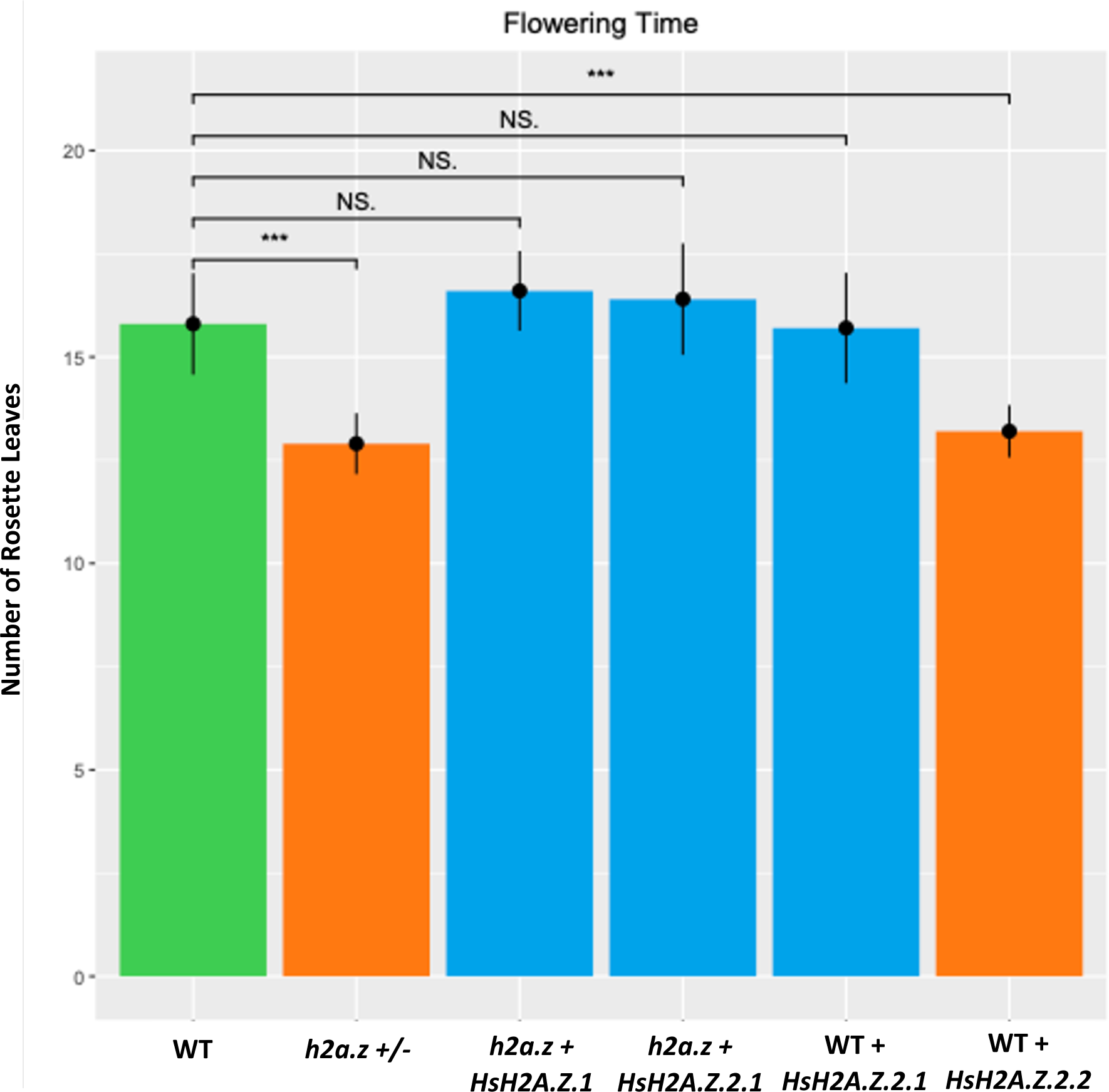
The average number of rosette leaves of WT plants, *h2a.z +/−*, *h2a.z + HsH2A.Z.1, h2a.z + HsH2A.Z.2.1*, WT + *HsH2A.Z.2.1*, and WT + *HsH2A.Z.2.2* transgenic plants at flowering. Flowering time (assayed as the average number of rosette leaves from 10 different plants per genotype at the time of bolting) is significantly different between WT and h2a.z +/− (with p value=1.305e-05), as well as between WT and WT + *HsH2A.Z.2.2* plants (with p value=4.204e-05). T-test was used for statistical analysis.

**S3 Figure.**
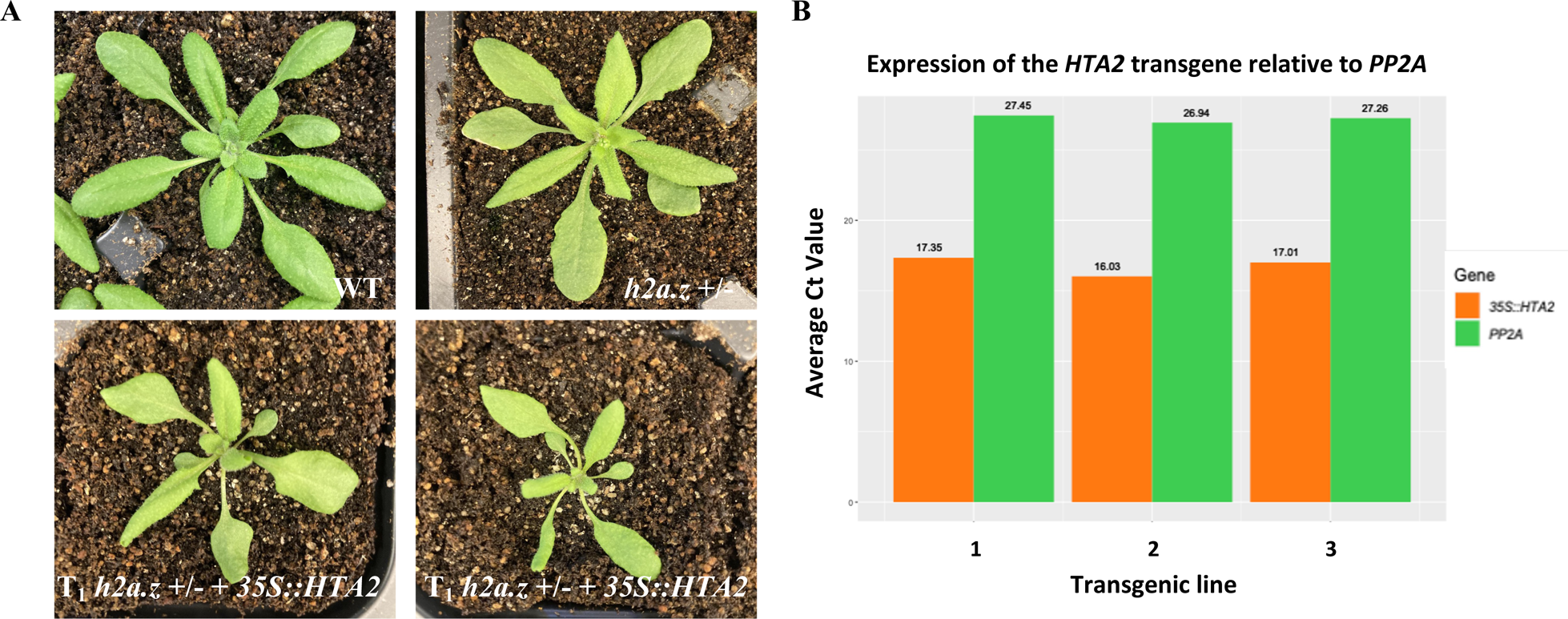
Overexpression of Arabidopsis canonical H2A histone HTA2 does not rescue *h2a.z +/−* phenotypic defects. **(A)** Two week old plants were grown under long-day conditions and individually photographed. **(B)** The cycle threshold (Ct) values of RT-qPCR assays of the *HTA2* transgene (orange bars) in three individual T_1_ plants (three biological replicates) relative to the Ct values of the endogenous control gene *PP2A* (green bars), as measured by RT-qPCR. Each biological replicate/transgenic plant had two technical RT-qPCR replicates.

**S4 Figure.**
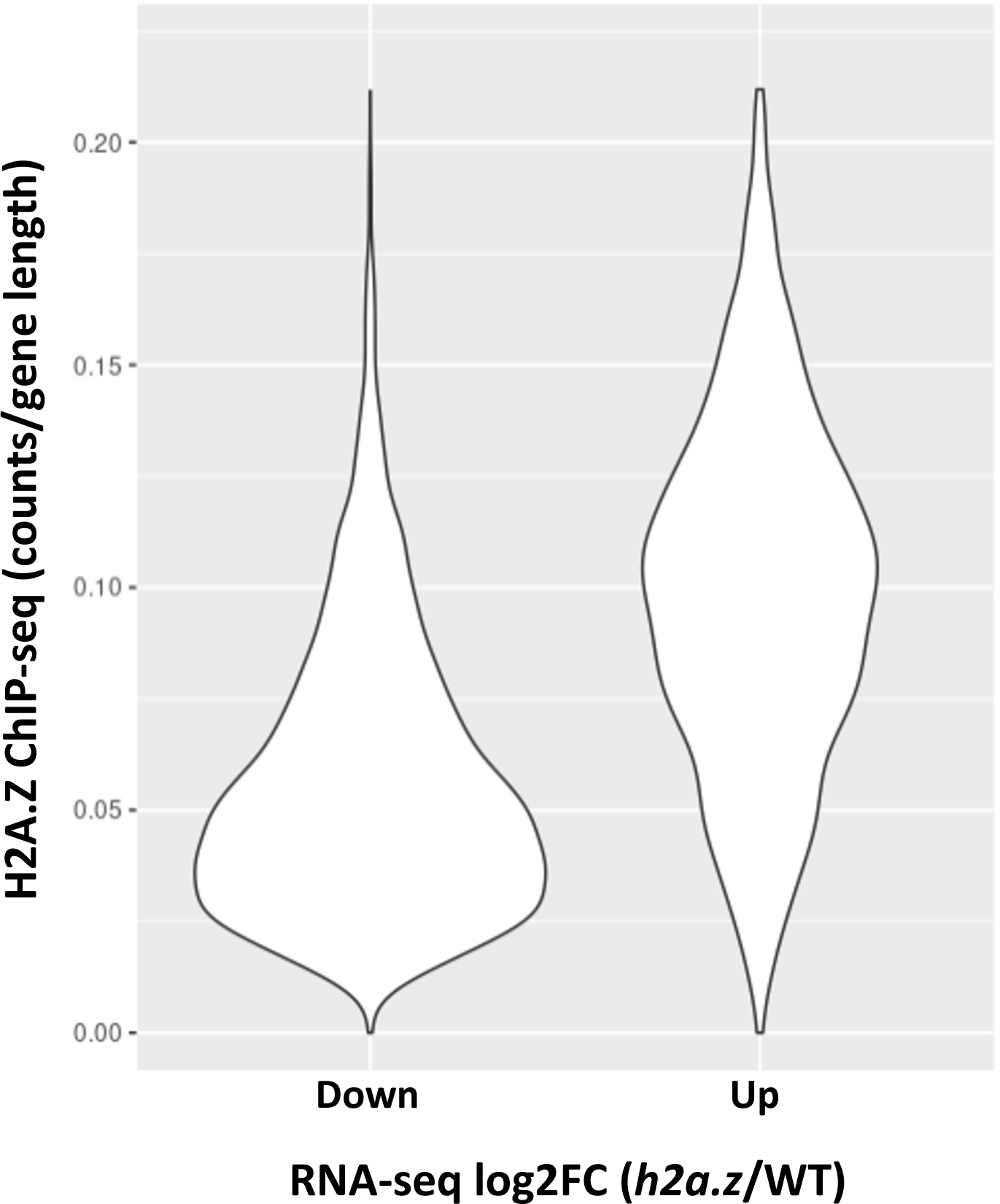
WT H2A.Z levels are higher at upregulated genes vs downregulated genes. Violin plots showing average H2A.Z enrichment in WT across the bodies of genes either downregulated (n = 3714) or upregulated (n =4685) in *h2a.z* plants. Counts are averaged over 3 DESeq2 normalized ChIP-seq replicates and corrected for gene length. Down and Up genes are defined as |L2FC| > 0.6 and padj < 0.05 with

**S5 Figure.**
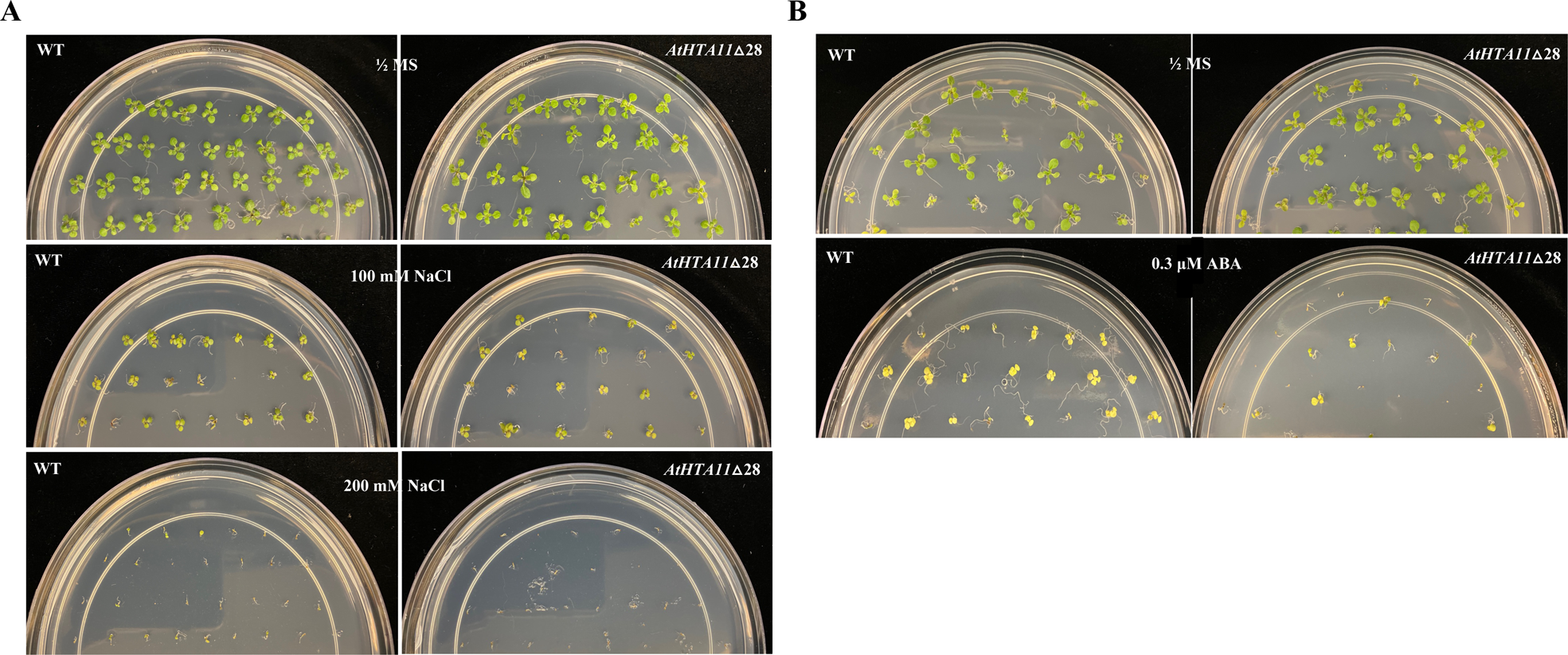
*AtHTA11△28* plants are more sensitive to high salt and ABA stresses than WT plants. Eleven-day-old seedlings grown on ½ MS agar plates or plates supplemented with either 100 mM or 200 mM NaCl **(A)** or supplemented with 0.3 μM Abscisic acid (ABA) **(B)**, were photographed and characterized for their ability to germinate and to produce green tissue. Each experiment was performed in duplicate

**S6 Figure.**
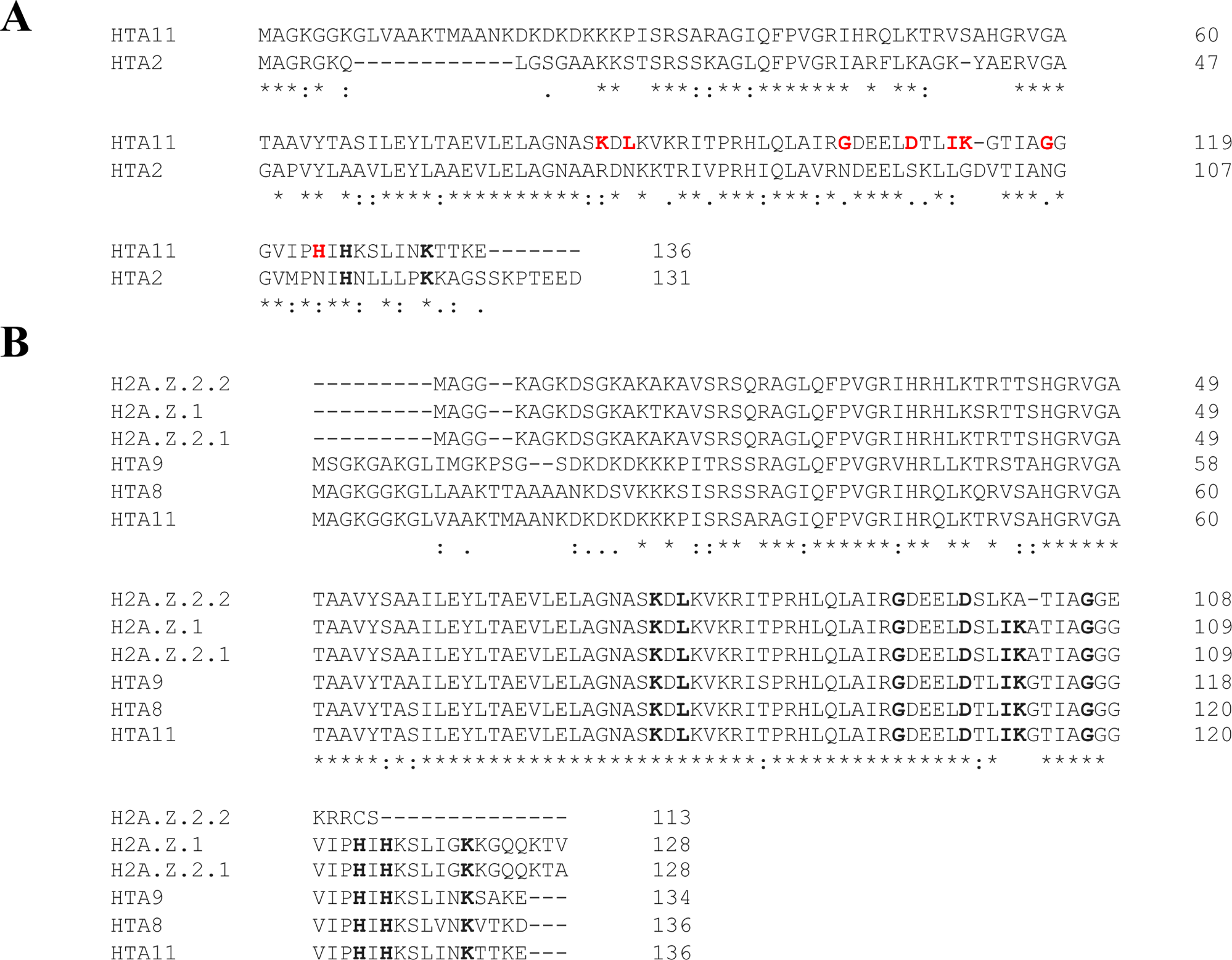
Conserved amino acids that contribute to H2A.Z unique function are found in both human and Arabidopsis H2A.Zs but not in Arabidopsis histone H2A. **(A)** Clustal Omega alignment between Arabidopsis HTA11 (H2A.Z) and HTA2 (core H2A) histones. Amino acids that are important for H2A.Z identity are highlighted in bold within the AtHTA11 sequence and amino acids exclusively found in AtHTA11 are highlighted in bold red and are not present in HTA2. **(B)** Clustal Omega alignment between human and Arabidopsis H2A.Zs. Amino acids that contribute to unique H2A.Z functions are highlighted in bold and are found in all Arabidopsis H2A.Zs and human H2A.Z.1 and H2A.Z.2.1, while in human H2A.Z.2.2. several key conserved residues at the C-terminal end are missing.

**S7 Figure.**
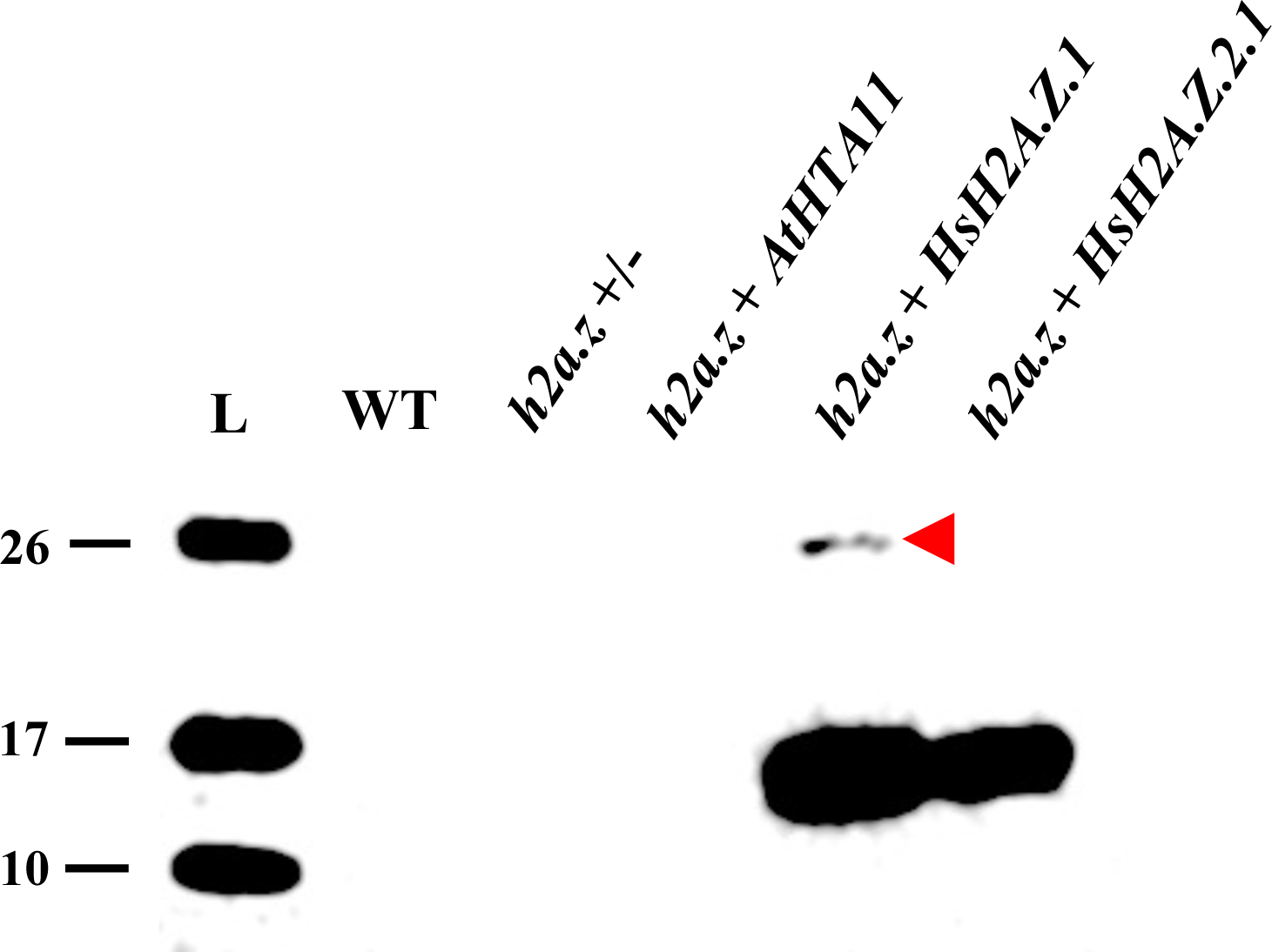
Human HsH2A.Z.1 appears to be monoubiquitinated when expressed in plants. The same western blot as shown in Figure 2C (middle panel, probed with an antibody against human H2A.Z) but overexposed, with the red arrowhead denoting the likely monoubiquitinated form of human HsH2A.Z.1 detected in the *h2a.z* +*HsH2A.Z.1* transgenic plants.

**S8 Figure.**
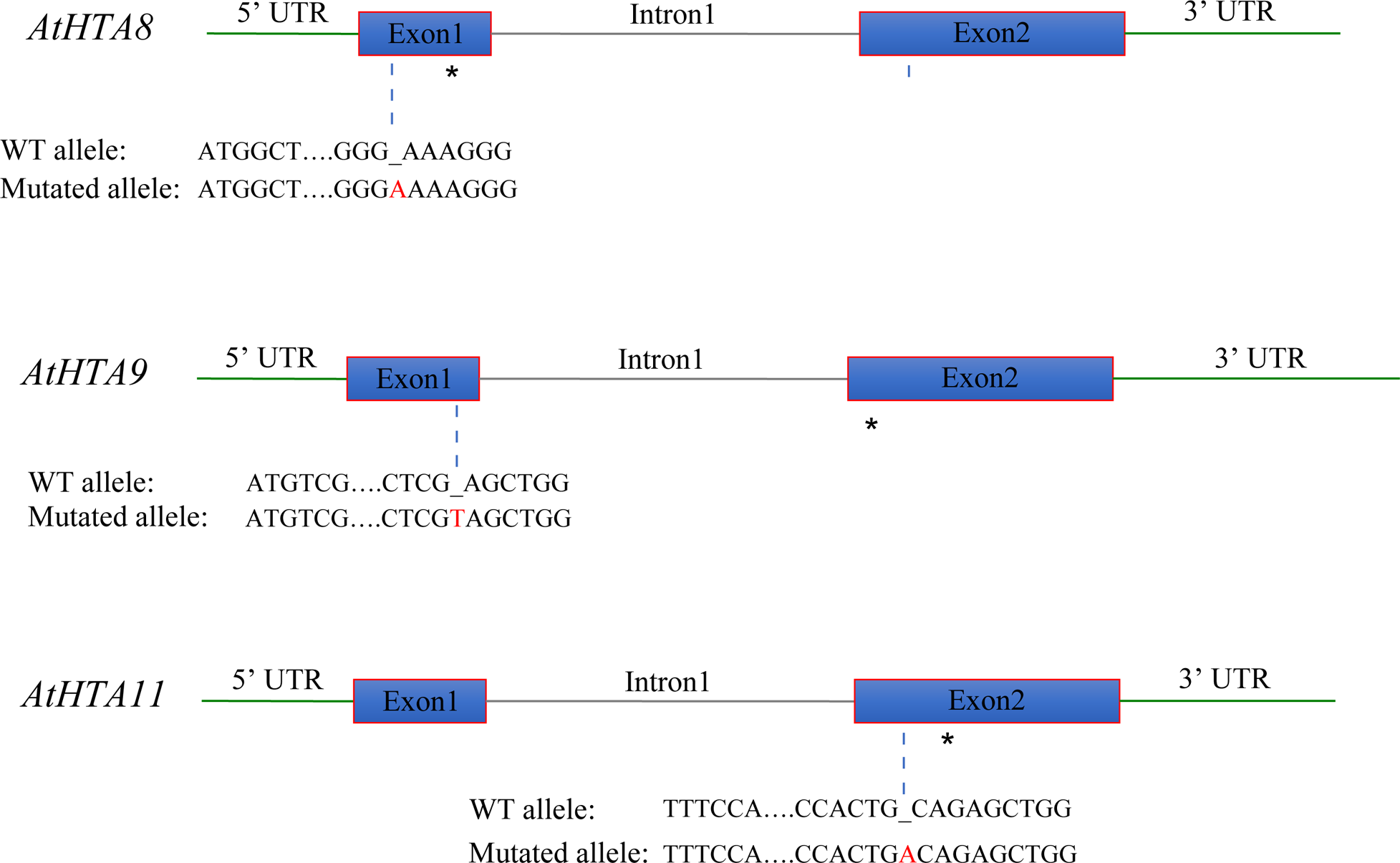
Graphical representation of Arabidopsis *h2a.z* CRISPR mutant alleles. For each H2A.Z gene the location and the type of CRISPR mutation is shown. All three genes have an addition of a single base pair causing a frame shift that leads to a premature stop codon (marked with an asterisk).

## REFERENCES

1. Talbert PB, Henikoff S. Histone variants at a glance. J Cell Sci. 2021;134(6).

2. Martire S, Banaszynski LA. The roles of histone variants in fine-tuning chromatin organization and function. Nat Rev Mol Cell Biol. 2020;21(9):522–41.

3. Foroozani M, Holder DH, Deal RB. Histone Variants in the Specialization of Plant Chromatin. Annu Rev Plant Biol. 2022;73:149–72.

4. Kreienbaum C, Paasche LW, Hake SB. H2A.Z’s ‘social’ network: functional partners of an enigmatic histone variant. Trends Biochem Sci. 2022;47(11):909–20.

5. Borg M, Jiang D, Berger F. Histone variants take center stage in shaping the epigenome. Curr Opin Plant Biol. 2021;61:101991.

6. Lewis TS, Sokolova V, Jung H, Ng H, Tan D. Structural basis of chromatin regulation by histone variant H2A.Z. Nucleic acids research. 2021;49(19):11379–91.

7. Bonisch C, Hake SB. Histone H2A variants in nucleosomes and chromatin: more or less stable? Nucleic acids research. 2012;40(21):10719–41.

8. Suto RK, Clarkson MJ, Tremethick DJ, Luger K. Crystal structure of a nucleosome core particle containing the variant histone H2A.Z. Nat Struct Biol. 2000;7(12):1121–4.

9. Xu Y, Ayrapetov MK, Xu C, Gursoy-Yuzugullu O, Hu Y, Price BD. Histone H2A.Z controls a critical chromatin remodeling step required for DNA double-strand break repair. Molecular cell. 2012;48(5):723–33.

10. Zhou BO, Wang SS, Xu LX, Meng FL, Xuan YJ, Duan YM, et al. SWR1 complex poises heterochromatin boundaries for antisilencing activity propagation. Molecular and cellular biology. 2010;30(10):2391–400.

11. Meneghini MD, Wu M, Madhani HD. Conserved histone variant H2A.Z protects euchromatin from the ectopic spread of silent heterochromatin. Cell. 2003;112(5):725–36.

12. Rosa M, Von Harder M, Cigliano RA, Schlogelhofer P, Mittelsten Scheid O. The Arabidopsis SWR1 chromatin-remodeling complex is important for DNA repair, somatic recombination, and meiosis. The Plant cell. 2013;25(6):1990–2001.

13. Coleman-Derr D, Zilberman D. Deposition of histone variant H2A.Z within gene bodies regulates responsive genes. PLoS genetics. 2012;8(10):e1002988.

14. March-Diaz R, Reyes J. The beauty of being a variant: H2A.Z and the SWR1 complex in plants. Molecular plant. 2009;2(4):565–77.

15. Giaimo BD, Ferrante F, Herchenrother A, Hake SB, Borggrefe T. The histone variant H2A.Z in gene regulation. Epigenetics Chromatin. 2019;12(1):37.

16. Colino-Sanguino Y, Clark SJ, Valdes-Mora F. The H2A.Z-nucleosome code in mammals: emerging functions. Trends Genet. 2022;38(5):516.

17. Faast R, Thonglairoam V, Schulz TC, Beall J, Wells JR, Taylor H, et al. Histone variant H2A.Z is required for early mammalian development. Current biology : CB. 2001;11(15):1183–7.

18. Lamaa A, Humbert J, Aguirrebengoa M, Cheng X, Nicolas E, Cote J, et al. Integrated analysis of H2A.Z isoforms function reveals a complex interplay in gene regulation. eLife. 2020;9.

19. Sales-Gil R, Kommer DC, de Castro IJ, Amin HA, Vinciotti V, Sisu C, et al. Non-redundant functions of H2A.Z.1 and H2A.Z.2 in chromosome segregation and cell cycle progression. EMBO Rep. 2021;22(11):e52061.

20. Bonisch C, Schneider K, Punzeler S, Wiedemann SM, Bielmeier C, Bocola M, et al. H2A.Z.2.2 is an alternatively spliced histone H2A.Z variant that causes severe nucleosome destabilization. Nucleic acids research. 2012;40(13):5951–64.

21. Wratting D, Thistlethwaite A, Harris M, Zeef LA, Millar CB. A conserved function for the H2A.Z C terminus. J Biol Chem. 2012;287(23):19148–57.

22. Deal R, Topp C, McKinney E, Meagher R. Repression of flowering in Arabidopsis requires activation of FLOWERING LOCUS C expression by the histone variant H2A.Z. The Plant cell. 2007;19(1):74–83.

23. Zilberman D, Coleman-Derr D, Ballinger T, Henikoff S. Histone H2A.Z and DNA methylation are mutually antagonistic chromatin marks. Nature. 2008;456(7218):125–9.

24. Choi K, Park C, Lee J, Oh M, Noh B, Lee I. *Arabidopsis* homologs of components of the SWR1 complex regulate flowering and plant development. Development (Cambridge, England). 2007;134(10):1931–41.

25. March-Diaz R, Garcia-Dominguez M, Lozano-Juste J, Leon J, Florencio F, Reyes J. Histone H2A.Z and homologues of components of the SWR1 complex are required to control immunity in Arabidopsis. The Plant journal : for cell and molecular biology. 2008;53(3):475–87.

26. Colino-Sanguino Y, Clark SJ, Valdes-Mora F. H2A.Z acetylation and transcription: ready, steady, go! Epigenomics. 2016;8(5):583–6.

27. Law C, Cheung P. Expression of Non-acetylatable H2A.Z in Myoblast Cells Blocks Myoblast Differentiation through Disruption of MyoD Expression. The Journal of biological chemistry. 2015;290(21):13234–49.

28. Bellucci L, Dalvai M, Kocanova S, Moutahir F, Bystricky K. Activation of p21 by HDAC inhibitors requires acetylation of H2A.Z. PLoS One. 2013;8(1):e54102.

29. Ku M, Jaffe JD, Koche RP, Rheinbay E, Endoh M, Koseki H, et al. H2A.Z landscapes and dual modifications in pluripotent and multipotent stem cells underlie complex genome regulatory functions. Genome Biol. 2012;13(10):R85.

30. Dalvai M, Fleury L, Bellucci L, Kocanova S, Bystricky K. TIP48/Reptin and H2A.Z requirement for initiating chromatin remodeling in estrogen-activated transcription. PLoS Genet. 2013;9(4):e1003387.

31. Binda O, Sevilla A, LeRoy G, Lemischka IR, Garcia BA, Richard S. SETD6 monomethylates H2AZ on lysine 7 and is required for the maintenance of embryonic stem cell self-renewal. Epigenetics. 2013;8(2):177–83.

32. Tsai CH, Chen YJ, Yu CJ, Tzeng SR, Wu IC, Kuo WH, et al. SMYD3-Mediated H2A.Z.1 Methylation Promotes Cell Cycle and Cancer Proliferation. Cancer Res. 2016;76(20):6043–53.

33. Nie WF, Lei M, Zhang M, Tang K, Huang H, Zhang C, et al. Histone acetylation recruits the SWR1 complex to regulate active DNA demethylation in Arabidopsis. Proceedings of the National Academy of Sciences of the United States of America. 2019;116(33):16641–50.

34. Jinek M, Chylinski K, Fonfara I, Hauer M, Doudna JA, Charpentier E. A programmable dual-RNA-guided DNA endonuclease in adaptive bacterial immunity. Science. 2012;337(6096):816–21.

35. Hartl M, Fussl M, Boersema PJ, Jost JO, Kramer K, Bakirbas A, et al. Lysine acetylome profiling uncovers novel histone deacetylase substrate proteins in Arabidopsis. Mol Syst Biol. 2017;13(10):949.

36. Brewis HT, Wang AY, Gaub A, Lau JJ, Stirling PC, Kobor MS. What makes a histone variant a variant: Changing H2A to become H2A.Z. PLoS genetics. 2021;17(12):e1009950.

37. Jiang D, Kong NC, Gu X, Li Z, He Y. Arabidopsis COMPASS-like complexes mediate histone H3 lysine-4 trimethylation to control floral transition and plant development. PLoS Genet. 2011;7(3):e1001330.

38. Benhamed M, Bertrand C, Servet C, Zhou DX. Arabidopsis GCN5, HD1, and TAF1/HAF2 interact to regulate histone acetylation required for light-responsive gene expression. Plant Cell. 2006;18(11):2893–903.

39. Bieluszewski T, Sura W, Dziegielewski W, Bieluszewska A, Lachance C, Kabza M, et al. NuA4 and H2A.Z control environmental responses and autotrophic growth in Arabidopsis. Nat Commun. 2022;13(1):277.

40. Crevillen P, Gomez-Zambrano A, Lopez JA, Vazquez J, Pineiro M, Jarillo JA. Arabidopsis YAF9 histone readers modulate flowering time through NuA4-complex-dependent H4 and H2A.Z histone acetylation at FLC chromatin. New Phytol. 2019;222(4):1893–908.

41. Murashige T, Skoog F. A revised medium for rapid growth and bio assays with tobacco tissue cultures. Physiologia Plantarum. 1962;15:473–97.

42. Clough SJ, Bent AF. Floral dip: a simplified method for Agrobacterium-mediated transformation of Arabidopsis thaliana. The Plant journal : for cell and molecular biology. 1998;16(6):735–43.

43. Wang ZP, Xing HL, Dong L, Zhang HY, Han CY, Wang XC, et al. Egg cell-specific promoter-controlled CRISPR/Cas9 efficiently generates homozygous mutants for multiple target genes in Arabidopsis in a single generation. Genome biology. 2015;16(1):144.

44. Curtis MD, Grossniklaus U. A gateway cloning vector set for high-throughput functional analysis of genes in planta. Plant Physiol. 2003;133(2):462–9.

45. Earley KW, Haag JR, Pontes O, Opper K, Juehne T, Song K, et al. Gateway-compatible vectors for plant functional genomics and proteomics. The Plant journal : for cell and molecular biology. 2006;45(4):616–29.

46. Czechowski T, Stitt M, Altmann T, Udvardi MK, Scheible WR. Genome-wide identification and testing of superior reference genes for transcript normalization in Arabidopsis. Plant physiology. 2005;139(1):5–17.

47. Luo YX, Hou XM, Zhang CJ, Tan LM, Shao CR, Lin RN, et al. A plant-specific SWR1 chromatin-remodeling complex couples histone H2A.Z deposition with nucleosome sliding. The EMBO journal. 2020;39(7):e102008.

48. Sijacic P, Holder DH, Bajic M, Deal RB. Methyl-CpG-binding domain 9 (MBD9) is required for H2A.Z incorporation into chromatin at a subset of H2A.Z-enriched regions in the Arabidopsis genome. PLoS genetics. 2019;15(8):e1008326.

49. Zhao L, Xie L, Zhang Q, Ouyang W, Deng L, Guan P, et al. Integrative analysis of reference epigenomes in 20 rice varieties. Nat Commun. 2020;11(1):2658.

50. https://www.bioinformatics.babraham.ac.uk/projects/trim_galore/.

51. Hou X, Wang D, Cheng Z, Wang Y, Jiao Y. A near-complete assembly of an Arabidopsis thaliana genome. Molecular plant. 2022;15(8):1247–50.

52. Langmead B, Salzberg SL. Fast gapped-read alignment with Bowtie 2. Nature methods. 2012;9(4):357–9.

53. Li H, Handsaker B, Wysoker A, Fennell T, Ruan J, Homer N, et al. The Sequence Alignment/Map format and SAMtools. Bioinformatics (Oxford, England). 2009;25(16):2078–9.

54. http://broadinstitute.github.io/picard/.

55. Quinlan AR, Hall IM. BEDTools: a flexible suite of utilities for comparing genomic features. Bioinformatics (Oxford, England). 2010;26(6):841–2.

56. Kent WJ, Zweig AS, Barber G, Hinrichs AS, Karolchik D. BigWig and BigBed: enabling browsing of large distributed datasets. Bioinformatics (Oxford, England). 2010;26(17):2204–7.

57. Ramirez F, Dundar F, Diehl S, Gruning BA, Manke T. deepTools: a flexible platform for exploring deep-sequencing data. Nucleic acids research. 2014;42(Web Server issue):W187–91.

58. Liao Y, Smyth GK, Shi W. featureCounts: an efficient general purpose program for assigning sequence reads to genomic features. Bioinformatics (Oxford, England). 2014;30(7):923–30.

59. Love MI, Huber W, Anders S. Moderated estimation of fold change and dispersion for RNA-seq data with DESeq2. Genome biology. 2014;15(12):550.

60. Wickham H. ggplot2: Elegant graphics for data analysis. Springer Verlag. 2016.

61. Dobin A, Davis CA, Schlesinger F, Drenkow J, Zaleski C, Jha S, et al. STAR: ultrafast universal RNA-seq aligner. Bioinformatics (Oxford, England). 2013;29(1):15–21.

